# Structural and biophysical characterisation of ubiquitin variants that specifically inhibit the ubiquitin conjugating enzyme Ube2d2

**DOI:** 10.1101/2024.03.10.583603

**Authors:** Jeffery M.R.B. McAlpine, Jingyi Zhu, Nicholas Pudjihartono, Joan Teyra, Michael J. Currie, Renwick C.J. Dobson, Sachdev S. Sidhu, Catherine L. Day, Adam J. Middleton

**Affiliations:** Department of Biochemistry, School of Biomedical Sciences, University of Otago, Dunedin, New Zealand; Department of Pharmacy, University of Waterloo, Kitchener, Ontario, Canada; Biomolecular Interaction Centre, Maurice Wilkins Centre for Biodiscovery, MacDiarmid Institute for Advanced Materials and Nanotechnology; and School of Biological Sciences, University of Canterbury, Christchurch 8140, New Zealand; Department of Biochemistry and Pharmacology, Bio21 Molecular Science and Biotechnology Institute, University of Melbourne, Parkville, Victoria, 3010, Australia

**Keywords:** ubiquitin, ubiquitin conjugating, E2 enzyme, inhibitor, protein-protein interaction

## Abstract

The ubiquitin conjugating E2 enzymes play a central role in ubiquitin transfer. Disruptions to the ubiquitin system are implicated in multiple diseases and as a result, molecules that modulate the activity of the ubiquitin system are of interest. E2 enzyme function is reliant on interactions with partner proteins and disruption of these is an effective way of modulating activity. Here, we report the discovery of ubiquitin variants (UbVs) that inhibit the E2 enzyme, Ube2d2 (UbcH5b). The six UbVs identified inhibit ubiquitin chain building, and structural and biophysical characterisation of two of these demonstrate they bind to Ube2d2 with low micromolar affinity and high specificity within the Ube2d family of E2 enzymes. Both characterised UbVs bind at a site that overlaps with E1 binding, while the more inhibitory UbV blocks a critical non-covalent ubiquitin binding site on the E2 enzyme. The discovery of novel protein-based ubiquitin derivatives that inhibit protein-protein interactions is an important step towards discovery of small molecules that inhibit the activity of E2 enzymes. Furthermore, the specificity of the UbVs within the Ube2d family suggests that it may be possible to develop tools to selectively inhibit highly related E2 enzymes.

## Introduction

The post-translational modification of proteins with ubiquitin is required for almost all aspects of eukaryotic cellular functions. Importantly, ubiquitin plays a central role in protein degradation where it regulates protein homeostasis and the cell cycle [1]. Ubiquitin also plays non-degradative roles and can regulate the timing of cell signalling and packaging of DNA. Consequently, disruptions to the post-translational modification of proteins with ubiquitin are associated with disease, including cancer and neurodegenerative diseases [2,3].

Attachment of ubiquitin to substrates relies on a three-enzyme cascade comprising a ubiquitin activating E1 enzyme, a ubiquitin conjugating E2 enzyme, and an E3 ligase. Ubiquitin usually modifies a primary amine on a substrate protein (either a lysine side chain or unmodified protein N terminus), but can also target threonine or serine side chains, as well as other unconventional targets [4]. The ubiquitin machinery can also produce ubiquitin chains of different types that have distinct consequences for the attached substrate protein [5]. In many cases, the E2 enzymes are the determinant of this code because they specify the exact linkage type of the ubiquitin chains. In the E2 family (∼40 in humans) some enzymes only produce one chain type, while others can synthesise disparate ubiquitin chains or add ubiquitin directly to substrates. This latter group includes a family of promiscuous E2 enzymes called the Ube2d family (also known as the UbcH5 family), which contains four highly-similar members, Ube2d1, Ube2d2, Ube2d3, and Ube2d4.

The Ube2d family plays important roles in cells, including mediating DNA repair processes [6], regulating the NF-κB/IκB pathway [7], and promoting degradation of p53 leading to inhibition of apoptosis [8]. The activity of the Ube2d family is reliant on its ability to interact with an E1 enzyme, diverse E3 ligases, and ubiquitin [9]. Importantly, the Ube2d family also binds ubiquitin non-covalently at an extra ubiquitin binding site, called the ‘backside’ site, that enhances ubiquitin chain building (processivity) [10]. This feature is partially due to an increase in the local concentration of ubiquitin at locations where the E2 enzymes are active, but it has also been shown that backside binding stabilises the Ube2d-E3 interaction to heighten ubiquitin transfer activity [11,12].

Generally, a hydrophobic surface on ubiquitin interacts with E2 enzymes and other components of the ubiquitin machinery. In 2013, a massive library of ubiquitin variants (UbVs) was developed where this surface of ubiquitin was diversified [13]. This library led to the discovery of specific and potent modulators of proteins in the ubiquitin system, including deubiquitinases [14], E3 ligases [15], ubiquitin interacting motifs [16], and E2 enzymes [17,18]. For the E2 enzymes, UbVs were identified that bound the backside of three E2 enzymes and their presence resulted in a decrease in ubiquitin transfer activity [17]. In a more recent example, UbVs were discovered that inhibited the activity of the E2, Ube2k, by sterically occluding the E2:E1 interface [18]. Together, these results highlight that disrupting protein-protein interactions with UbVs is an effective means for modulating the activity of E2 enzymes.

Here, we have used phage display to discover UbVs that bind to Ube2d2 at sites distinct from the backside by using a variant of the E2 enzyme that disrupts backside ubiquitin binding. We report the discovery of six UbVs that reduce the formation of ubiquitin chains, with two of these being the focus of further study. Crystallographic and biophysical analysis demonstrate that the UbVs bind to a highly-similar site on Ube2d2 that likely disrupts interaction with the E1 ubiquitin activating enzyme. However, the more inhibitory UbV binds as a dimer, and its additional binding site on Ube2d2 overlaps with the backside binding, resulting in an enhanced ability to inhibit ubiquitin chain formation. We also show that the UbVs can distinguish members of the Ube2d family of E2 enzymes, suggesting that there are opportunities to discover compounds that specifically impede the activity of distinct E2 enzymes.

## Results

### Selection against a Ube2d2 mutant

An earlier study identified UbVs that bind to the backside of the E2 enzymes Ube2d1, Ube2v1 and Ube2g1, and these UbVs overlap very well with the allosteric binding site of ubiquitin [17]. Here, we wanted to discover UbVs that can modulate the activity of Ube2d2 by binding to other sites, and therefore we used a mutated form of Ube2d2, Ube2d2^S22R^, which contains a mutation that disrupts backside ubiquitin binding [10,19]. Recombinant Ube2d2^S22R^ was expressed in *Escherichia coli* with a C-terminal Avi tag to allow it to be biotinylated for indirect immobilisation to plates. After expression, purification, and immobilisation of Ube2d2^S22R^, selection was performed using phage display with a library containing approximately 10^9^ UbVs. This UbV library is identical to the one described earlier [18]. Briefly, the library contains synthetic constructs based on ubiquitin from *Homo sapiens* that have been diversified on the hydrophobic surfaces around Ile44, as well as along the C-terminal β-strand and last seven residues of ubiquitin. The UbV library was produced using a soft-randomisation strategy to ensure there was a preference for wild-type amino acids so that the fold was not disrupted [13]. Four rounds of selection using Ube2d2^S22R^ were performed with the UbV library, and binding of UbVs was confirmed by ELISAs. This resulted in the selection of six UbVs (UbV.d2.1–UbV.d2.6, hereafter referred to as UbV.1–UbV.6, Fig 1A) that were amplified including an N-terminal FLAG tag and cloned into a vector that encoded an N-terminal cleavable His tag. Subsequently, the UbVs were expressed in *E. coli* and purified.

**Figure 1.**
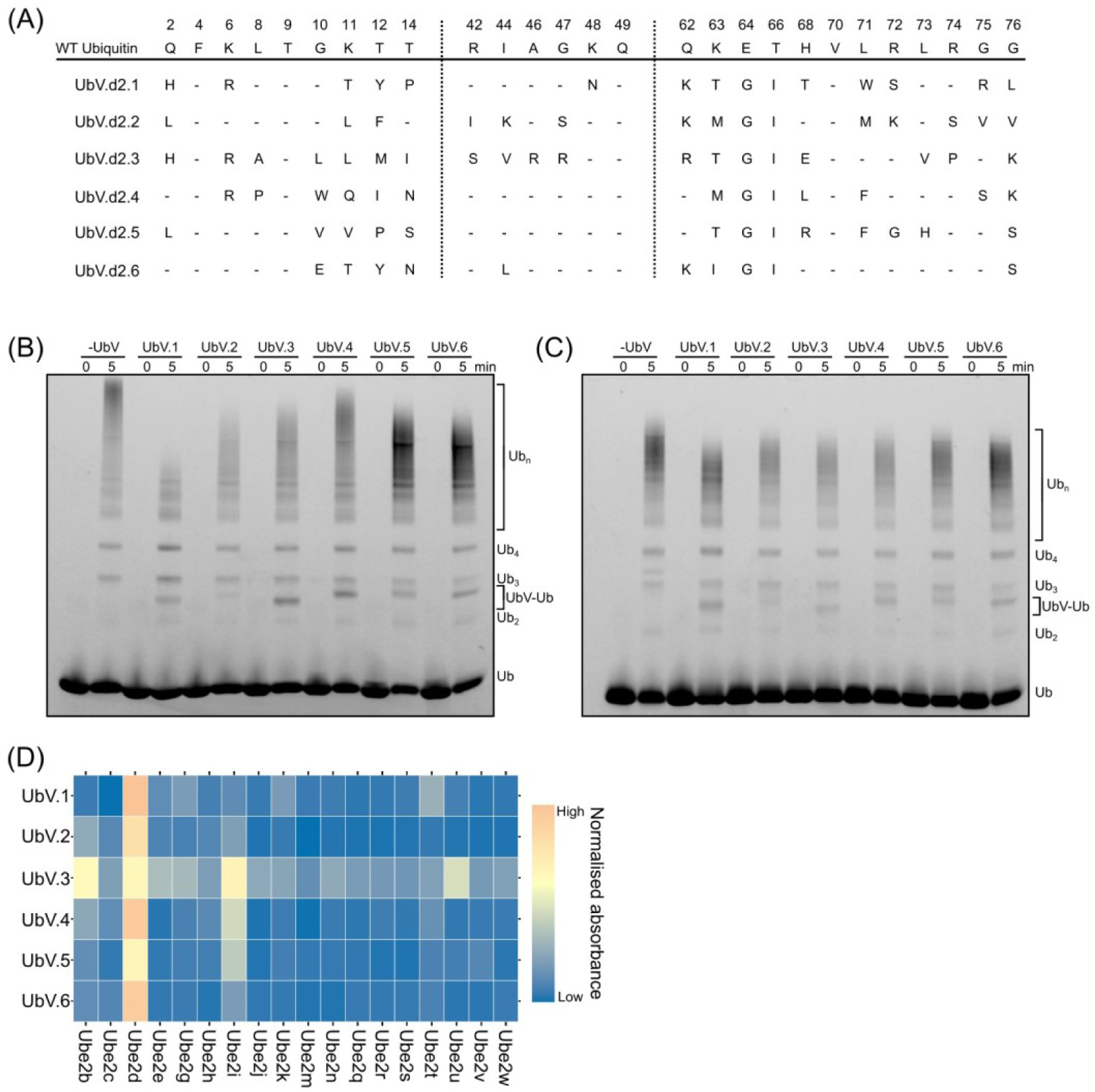
Identification of six UbVs that inhibit ubiquitin chain formation by Ube2d2. **(A)** Outline of the UbVs identified with the positions diversified in the library shown below the wild-type ubiquitin sequence. Wild-type amino acids are indicated with a hyphen. **(B, C)** Multi-turnover chain-building assay in the presence and absence of the six UbVs labelled UbV.1–UbV.6 with (B) Ube2d2 and **(C)** Ube2d2^S22R^. Ub refers to ubiquitin. Cy3-labelled ubiquitin imaged at 600 nm. **(D)** Specificity screen against a representative panel of E2 enzymes where the colours represent the normalised signal from ELISA experiments (mean of four technical replicates).

### The UbVs inhibit ubiquitin chain-building

In order to assess whether the isolated UbVs modulate ubiquitin transfer, we used a ubiquitin chain-building assay. For this assay, E1 ubiquitin activating enzyme, wild-type Ube2d2, and the RING domain of an E3 ligase (RNF12) were mixed with fluorescently-labelled ubiquitin and incubated at 37 °C. The extent of chain formation was assessed by SDS-PAGE. In the absence of the UbVs, extensive high molecular weight ubiquitin chains are apparent, while in the presence of each UbV, the extent of longer chain formation is decreased (Fig 1B). The decrease in long chains was particularly evident for UbV.1. To assess whether the UbVs could inhibit Ube2d2^S22R^, another chain-building assay was performed with the E2 variant. Interestingly, with Ube2d2^S22R^ all the UbVs resulted in slight decreases in the length of ubiquitin chains (Fig 1C), but none of the UbVs disrupted the activity of Ube2d2^S22R^ to the same extent as for the wild-type E2 enzyme.

Selection technology is a powerful tool for discovering tight-binding and highly-specific binders for particular protein targets [20]. However, E2 enzymes contain a conserved core domain, and many E2 enzymes are very similar. To assess whether the UbVs selected were specific for Ube2d2, we conducted an ELISA experiment where we immobilised a representative selection of eighteen E2 enzymes to a 96-well plate [18]. Analysis of binding showed that all the UbVs bound to Ube2d2, and most were specific (Fig 1D). Together, these experiments revealed the discovery of a set of UbVs that selectively inhibit the activity of wild-type and an S22R variant of Ube2d2, with UbV.1 being the most potent.

### Crystal structures of UbVs with Ube2d2 and Ube2d2^S22R^

To understand how the UbVs inhibit the chain-building activity of Ube2d2, we attempted to crystallise the UbVs in complex with Ube2d2. Of the six UbVs isolated, crystals were obtained for the UbV.1-Ube2d2 complex and the UbV.3-Ube2d2^S22R^ complex. Datasets were collected at the MX2 beamline at the Australian Synchrotron and their structures were solved by molecular replacement using models of Ube2d2 (PDB 3TGD) and wild-type ubiquitin (PDB 1UBQ) (Table 1).

**Table 1.** Crystallographic data collection and refinement statistics.

|  | UbV.d2.1 | UbV.d2.3 |
| --- | --- | --- |
| **PDB Entry** | 8VK8<sup>a</sup> | 8VK9 |
| **Data Collection** |  |  |
| Wavelength (Å) | 0.9537 | 0.9537 |
| Beamline | Australian synchrotron MX2 | Australian synchrotron MX2 |
| Resolution range | 41.0-3.0 (3.10-3.0)<sup>b</sup> | 41.2-2.0 (2.07-2.0) |
| Space group | P3<sub>1</sub>21 | P1 |
| Unit cell dimensions |  |  |
| a, b, c (Å) | 129.77 129.77 123.11 | 55.85 56.31 117.50 |
| α, β, γ (°) | 90 90 120 | 86.49 78.47 60.32 |
| Total no. of reflections | 332055 (32417) | 243317 (25030) |
| Unique reflections | 24503 (2364) | 80298 (7996) |
| Multiplicity | 13.6 (13.7) | 3.0 (3.1) |
| Completeness (%) | 99.8 (98.6) | 97.5 (96.7) |
| R<sub>merge</sub> | 0.184 (2.00) | 0.0528 (0.765) |
| I/σ (I) | 17.6 (1.4) | 10.2 (1.2) |
| CC<sub>1/2</sub> | 0.999 (0.629) | 0.998 (0.678) |
| **Refinement** |  |  |
| No. of reflections | 24495 (2363) | 80211 (7976) |
| No. of reflections (free) | 1145 (93) | 4049 (421) |
| R<sub>work</sub> (%) | 0.247 (0.405) | 0.190 (0.333) |
| R<sub>free</sub> (%) | 0.294 (0.436) | 0.224 (0.378) |
| No. of atoms | 6499 | 7637 |
| Protein | 6499 | 7245 |
| Solvent | 0 | 362 |
| Ligand | 0 | 5 |
| Average B factor (Å<sup>2</sup>) | 107.6 | 63.7 |
| RMSD bonds (Å) | 0.002 | 0.003 |
| RMSD angles (°) | 0.54 | 0.71 |
| Ramachandran plot |  |  |
| Favoured (%) | 98.2 | 97.4 |
| Allowed (%) | 1.8 | 2.5 |
| Outliers (%) | 0 | 0.1 |

<sup>a</sup>Each structure was determined from a single crystal <sup>b</sup>Values for the highest-resolution shell are shown in parentheses

The structure of the more highly inhibitory UbV.1 with Ube2d2 was solved at 3.0 Å in the P3_1_21 spacegroup. Its asymmetric unit contains three Ube2d2 molecules, two of which each bind two UbV molecules (referred to as UbV.1a and UbV.1b, below). For the third Ube2d2 molecule, only one UbV (UbV.1a) was modelled. While there is positive density in the mFo-DFc electron density map at the expected position next to the third Ube2d2 molecule (chain G), no molecule could be reliably placed (Supp Fig 1). The interaction of UbV.1a with Ube2d2 is reliant on a hydrophobic core comprising Phe3, Tyr12, and Ile66 from UbV.1a. The interface is centred on Met30 along helix 1 of Ube2d2 and abuts its β1-β2 loop (Fig 2A,B). UbV.1b interacts with Ube2d2 along the opposing side of the β1-β2 loop of Ube2d2. Notably, the side chains of Ile44 and Val70 from UbV.1b pack against Phe51 from Ube2d2 (Fig 2C). Both of the UbV-Ube2d2 interactions bury a similar area (UbV.1a and UbV.1b bury 476 and 562 Å^2^, respectively) and are heavily reliant on the diversified residues from the UbV (Figure 2D).

**Figure 2.**
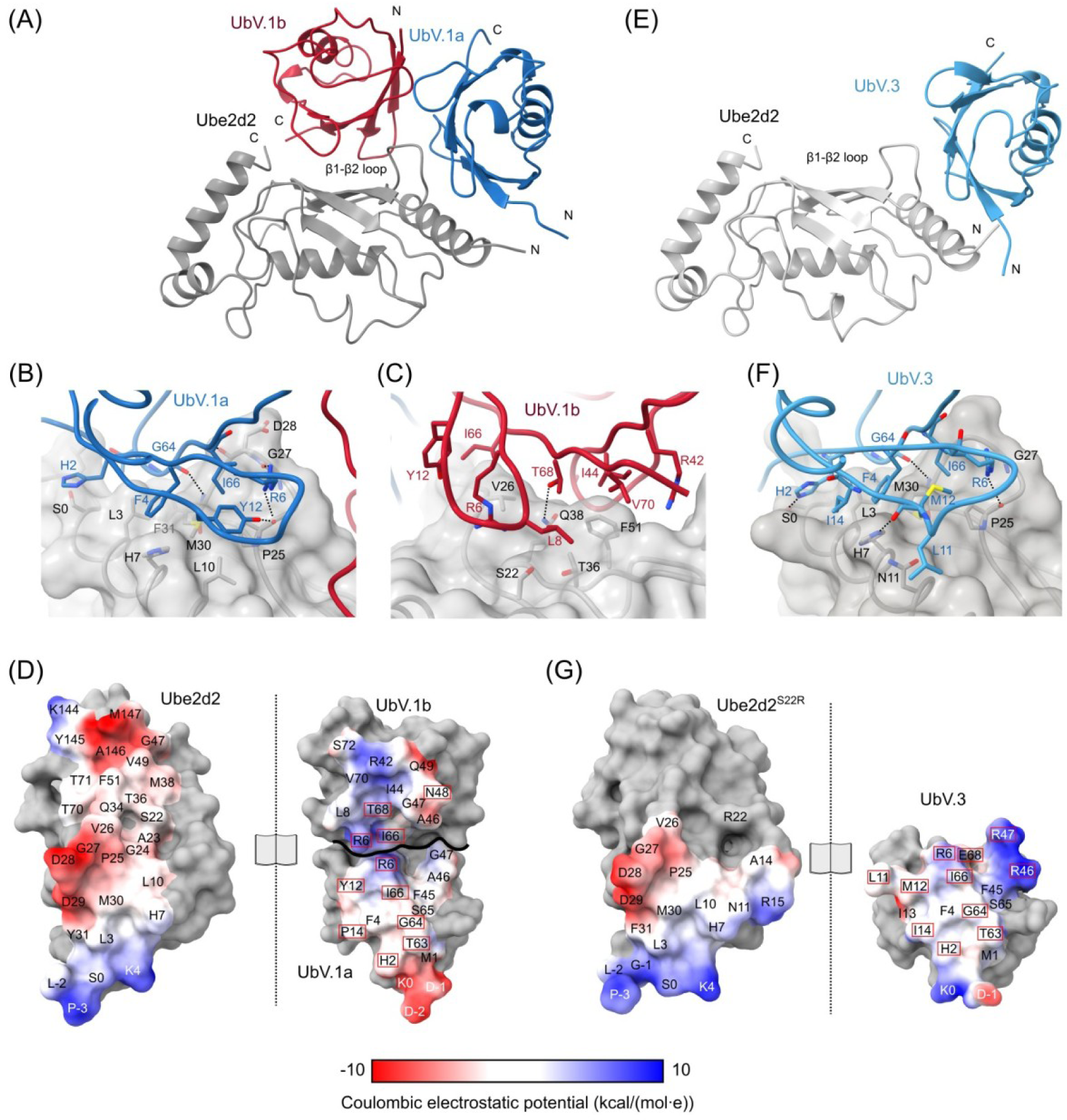
Crystal structure of two UbVs in complex with Ube2d2 and Ube2d2^S22R^. **(A)** Ribbon representation of the crystal structure of UbV.1 in complex with Ube2d2. The E2 enzyme is in grey while the two UbV.1 molecules are in blue and red. N and C termini are indicated. **(B,C)** Close up view of contacting residues shown as sticks from **(B)** UbV.1a and **(C)** UbV.1b with Ube2d2. Ube2d2 is shown as a semi-transparent surface. Red, blue, and yellow represent oxygen, nitrogen, and sulfur atoms, respectively. **(D)** Open-book surface representation showing the contacting residues between Ube2d2 and the two bound UbVs. Colouring is based off the Coulombic electrostatic potential of the proteins as calculated by Chimerax v1.6.1 [41]. Residues in the UbV that are diversified from wild-type ubiquitin are highlighted with red boxes. **(E,F,G)** Identical representation as for panels A-D but for the structure of UbV.3 with Ube2d2^S22R^.

While each of the two UbVs makes extensive contacts with Ube2d2, they also interact with each other (Supp Fig 2A). The UbV.1a-UbV.1b interface buries approximately 350 Å^2^ and is entirely hydrophobic with no polar contacts or predicted hydrogen bonds (Supp Fig 2B). Curiously, UbV.1a uses largely wild-type ubiquitin residues as part of its interface with its partner UbV, while UbV.1b is more reliant on diversified residues. The small nature of this interface makes it unlikely that UbV.1 forms a dimer in solution on its own and instead is only observed when the UbV is bound to the E2 enzyme.

The crystal structure of UbV.3 with Ube2d2^S22R^ was solved at 2.0 Å in the P1 spacegroup. The asymmetric unit contains four Ube2d2^S22R^ molecules each in complex with one molecule of UbV.3. The interaction between the UbV and Ube2d2^S22R^ is very similar to that observed for UbV.1a-Ube2d2, though UbV.1a is rotated by approximately 10° relative to UbV.3 (Supp Fig 2C). The interface buries approximately 700 Å^2^ and, similarly to the UbV.1-Ube2d2 interaction, is highly reliant on diversified residues (Fig 2E-G).

### UbV.1 and UbV.3 binding to Ube2d2 disrupt critical protein-protein interactions

To build ubiquitin chains on substrate proteins, E2 enzymes must first be charged with ubiquitin to form an E2∼Ubiquitin (E2∼Ub) conjugate. Charging of an E2 relies on its interaction with an E1 enzyme that is loaded with an activated ubiquitin molecule. Once the E2∼Ub conjugate is formed, it then pairs with an E3 ligase to catalyse transfer of ubiquitin to a substrate. Because disrupting any of these interactions would negatively affect activity, we investigated each in turn (Fig 3A-D). First, we overlayed both UbV-Ube2d2 structures with an E1-engaged E2 enzyme [21]. This overlay shows a substantial overlap in the binding site of the UbVs with part of the E1 enzyme interface on Ube2d (Fig 3A, pink surface). To establish whether the UbVs can disrupt engagement with an E1 enzyme, we performed E2 charging assays. For this experiment, a mixture of Ube2d2, E1, fluorescently-tagged ubiquitin, and ATP were incubated at 30 °C for 10 minutes with or without the two UbVs. The level of Ube2d2∼Ub conjugate was then detected by SDS-PAGE (Fig 3E). When the UbVs were in excess, we observed a decrease in the amount of charging, indicating that the UbVs block engagement of Ube2d2 with the E1 enzyme.

**Figure 3.**
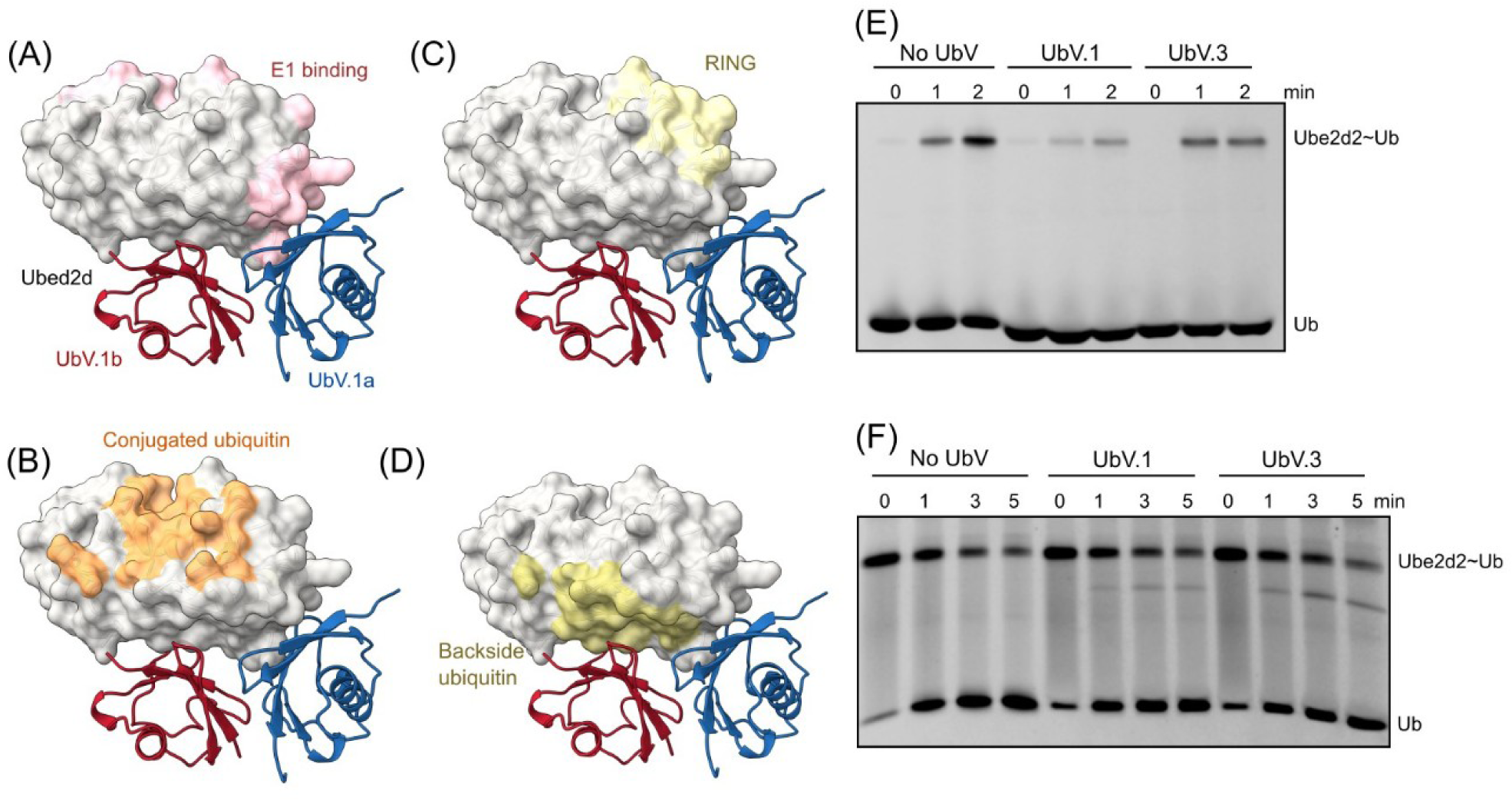
Functional analysis of the UbV-Ube2d2 complexes. **(A-D)** A surface representation of Ube2d2 in grey with UbV.1a in blue and UbV.1b in red. Panels indicate the interface with: **(A)** the E1 enzyme in pink; **(B)** the RING interface in yellow; **(C)** conjugated ubiquitin in orange; and **(D)** backside ubiquitin in yellow. UbV.3 is not shown for clarity. **(E)** A single-turnover E2 charging assay showing the formation of the Ube2d2∼Ubiquitin conjugate with or without the two UbVs. **(F)** A single-turnover E3-catalysed ubiquitin discharge of Ube2d2∼Ubiquitin conjugates with or without the UbVs. Both gels imaged at 600 nm.

Next we overlayed the structures with a Ube2d2∼Ub conjugate engaged with a RING domain [11]. This overlay shows that the UbVs bind adjacent to the RING and conjugated ubiquitin binding site (Fig 3B,C). To investigate if the UbVs impede interaction with the E3 ligase, we performed E3-catalysed ubiquitin discharge experiments using purified Ube2d2∼Ub conjugates. For these experiments, the RNF12^RING^ was used as an E3 ligase, and ubiquitin discharge was monitored with or without the UbVs. The level of ubiquitin discharge from Ube2d2 was assessed using SDS-PAGE (Fig 3F). This experiment showed that neither UbV affected E3-catalysed discharge. A subsequent pulldown experiment also confirmed that in the presence of the UbVs, there was no disruption to E3 binding (Supp Fig 3A).

Finally, an overlay of the UbV.1-Ube2d2 structure with Ube2d2 in complex with backside-bound ubiquitin revealed that UbV.1b appears to overlap with the allosteric binding site of ubiquitin (Fig 3D). This potential disruption of backside binding may explain why we observed more inhibition of ubiquitin chain formation by UbV.1. Altogether, these experiments suggest that UbV.1 and UbV.3 inhibit ubiquitin charging of Ube2d2 by inhibiting binding of the E1 enzyme, while UbV.1 likely also inhibits backside ubiquitin binding. These experiments also confirm that the UbVs do not have an effect on E3-catalyzed discharge.

### UbV.1 and UbV.3 form stable complexes with Ube2d2 and Ube2d2^S22R^

The crystal structures of UbV.1 with Ube2d2 and UbV.3 with Ube2d2^S22R^ suggest that they form stable complexes, but with different stoichiometry. To determine the affinity of the two UbVs for Ube2d2 and Ube2d2^S22R^, we used isothermal titration calorimetry (ITC). For this experiment we titrated UbV.1 and UbV.3 at approximately 200 µM into the E2 enzymes at around 20 µM (Fig 4). ITC confirmed that each of the UbVs formed tight complexes with Ube2d2 and Ube2d2^S22R^ with low micromolar binding constants. No binding between wild-type ubiquitin and Ube2d2 was detected (Supp Fig 3B).

**Figure 4.**
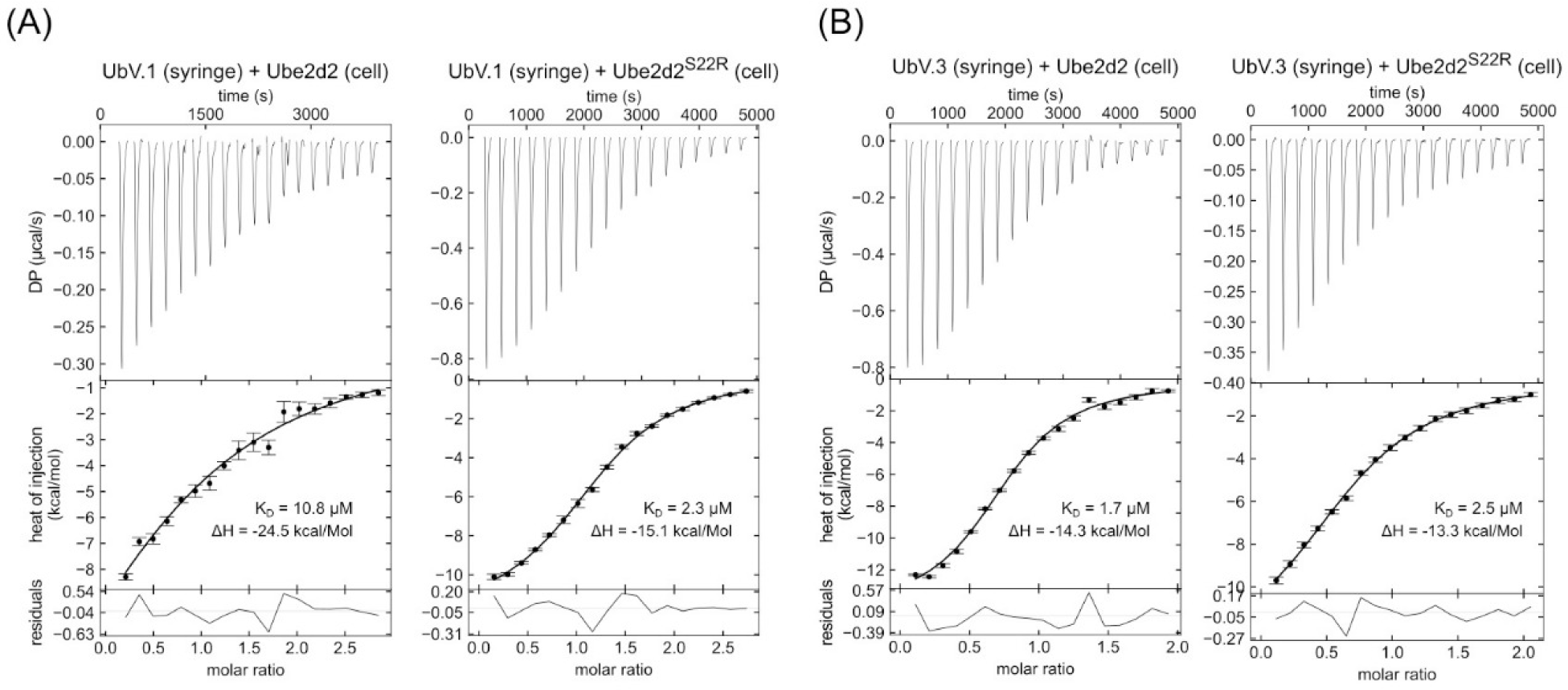
Measurement of UbV-Ube2d2 interactions. **(A)** ITC titrations of UbV.1 into Ube2d2 and Ube2d2^S22R^; and **(B)** of UbV.3 into Ube2d2 and Ube2d2^S22R^. The binding constant (K_D_) and the enthalpy change (ΔH) are shown.

We next measured the stoichiometry of the UbV-Ube2d2 complexes using analytical size-exclusion chromatography and analytical ultracentrifugation. For the size-exclusion chromatography experiments, Ube2d2 and Ube2d2^S22R^ were injected over an analytical S75 column followed by mixtures of the UbVs with either wild-type (Fig 5A) or the S22R variant of the E2 enzyme (Fig 5B). The complexes were prepared at a 2:1 ratio (100 µM of UbVs to 50 µM) to account for the possibility of different binding ratios as suggested by the crystal structure of UbV.1-Ube2d2. All four of the mixtures resulted in clear peak shifts, providing additional evidence that the UbVs form stable complexes with Ube2d2 (Fig 5A,B). Estimations of the molecular weight of the complexes by comparison with protein standards suggest that UbV.1 binds at a 2:1 ratio to Ube2d2, while UbV.3 binds at a 1:1 ratio. For the UbV-Ube2d2^S22R^ complexes, UbV.1 appears to elute between a 2:1 and a 1:1 ratio, suggesting that the S22R mutation might partially disrupt binding of one of the two UbVs. For the UbV.3 complex with Ube2d2^S22R^, a 1:1 ratio was observed.

**Figure 5.**
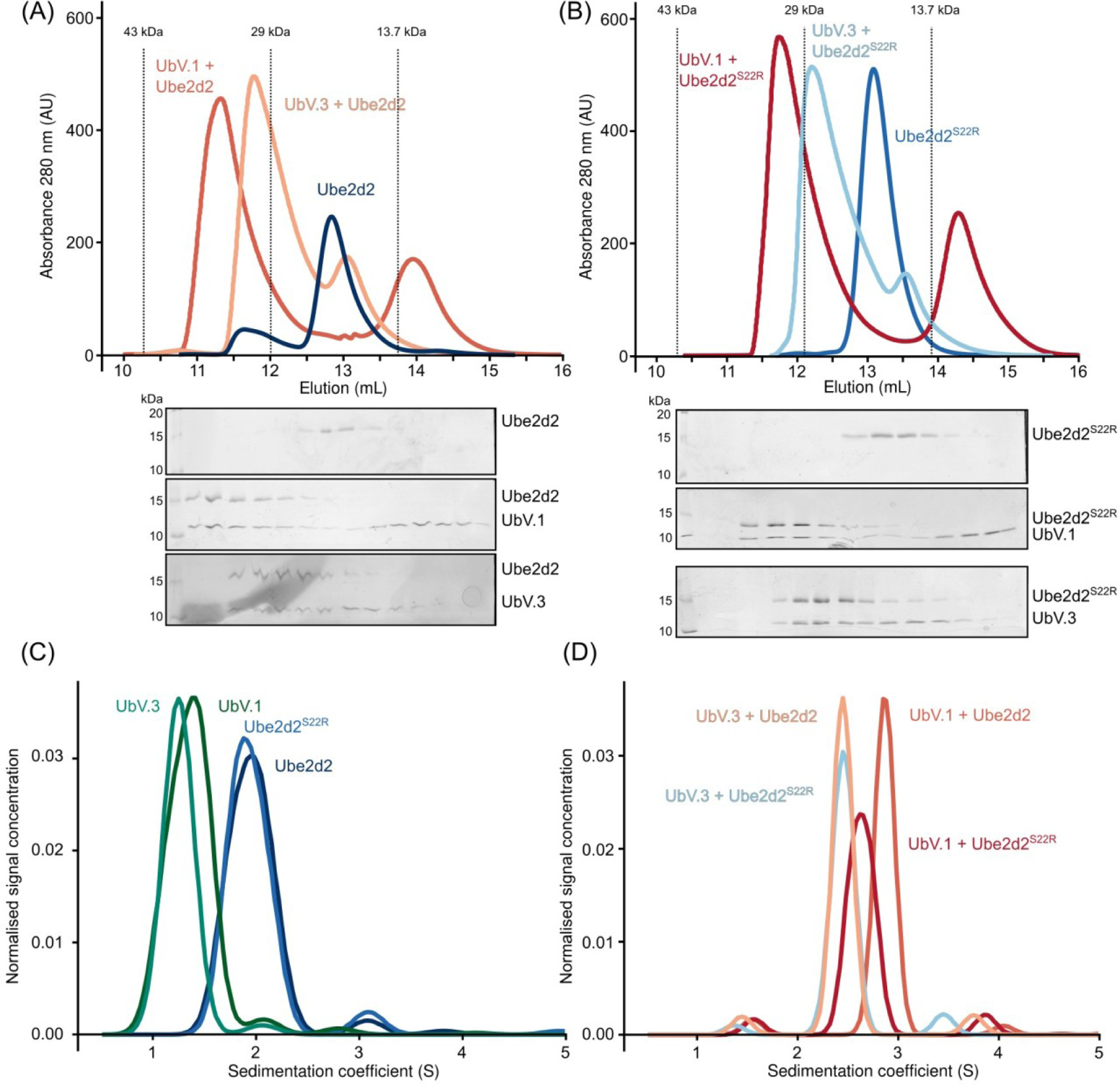
Determining the in solution stoichiometry of the UbV-Ube2d2 complexes. **(A, B)** Size-exclusion chromatography of UbV.1 and UbV.3 with **(A)** Ube2d2 and **(b)** Ube2d2^S22R^. Below shows the corresponding fractions. Formation of stable complexes is indicated by elution peaks shifting to the left. Protein standards were used to determine the molecular weights indicated with dotted lines. **(C)** Sedimentation velocity analysis of Ube2d2, Ube2d2^S22R^, UbV.1, and UbV.3 alone (detected at 280 nm). **(D)** Sedimentation velocity analysis of UbV-Ube2d2 complexes where Ube2d2 and Ube2d2^S22R^ were labelled with FITC and sedimentation tracked using the absorbance of FITC at 493 nm. As a result, only Ube2d2 and Ube2d2^S22R^ can be observed. Stable complexes are indicated by peak shifts to the right relative to panel C.

Analytical ultracentrifugation experiments using the sedimentation velocity method showed consistent results. When each protein was analysed alone (Fig 5C), all are largely monomeric with the expected masses: UbV.1 and UbV.3 are smaller with S values at ∼1.2 S (∼11 kDa), while Ube2d2 and Ube2d2^S22R^ have S values at ∼2 S (∼18 kDa). When Ube2d2 was mixed and incubated with UbV.3 or UbV.1 (Fig 5D orange traces), the distribution of Ube2d2 shifts to the right relative to Ube2d2 alone (UbV.3 to ∼2.3 S and UbV.1 to ∼2.9 S) consistent with binding of one UbV.3 and two UbV.1 molecules, as seen in the crystal structures (Fig 2A,E). Similarly, when Ube2d2^S22R^ was mixed and incubated with UbV.3 (Fig 5D, blue trace), the peak for Ube2d2^S22R^ shifts to ∼2.3 S, consistent with one UbV.3 molecule binding to Ube2d2^S22R^. However, when Ube2d2^S22R^ was mixed and incubated with UbV.1 (Fig 5D, red trace), the peak shifts right to a lesser extent, suggesting that one or two UbV.1 molecules can bind to Ube2d2^S22R^. These results are consistent with the analytical size-exclusion chromatography experiment (Fig 5B, red trace) and confirm that the UbVs form a stable complex with Ube2d2 and Ube2d2^S22R^ with differing stoichiometry.

### Specificity of the UbVs for E2 enzymes and the Ube2d family

The four members of the Ube2d family of E2 enzymes are highly conserved, with a sequence identity of greater than 85%, and they have largely been thought of as functionally equivalent (Supp Fig 4). In the crystal structures, UbV.1a and UbV.3 both make contacts with Ube2d2 that include amino acids that vary between members of the Ube2d family (Fig 6A, Supp Fig 4), suggesting that the UbVs could be specific for particular family members. To determine specificity, we prepared GST-fusion proteins of the two UbVs and performed pulldowns against the four members of the Ube2d family (Fig 6B). As expected, UbV.1 bound to Ube2d2, while it also interacted with Ube2d4. In contrast, UbV.3 only interacted with Ube2d2. Subsequent chain-building assays confirmed that UbV.1 could attenuate the activity of Ube2d2 as well as Ube2d4, but had no effect on Ube2d1 or Ube2d3 (Fig 6C). UbV.3 resulted in a slight decrease in chain formation with Ube2d3, while it inhibited the activity of Ube2d2 at a similar level as shown in Fig 1. Together with the specificity ELISAs (Fig 1D), these experiments suggest that UbV.1 binds only to Ube2d2 and Ube2d4, while UbV.3 is specific for Ube2d2 within the Ube2d family.

**Figure 6.**
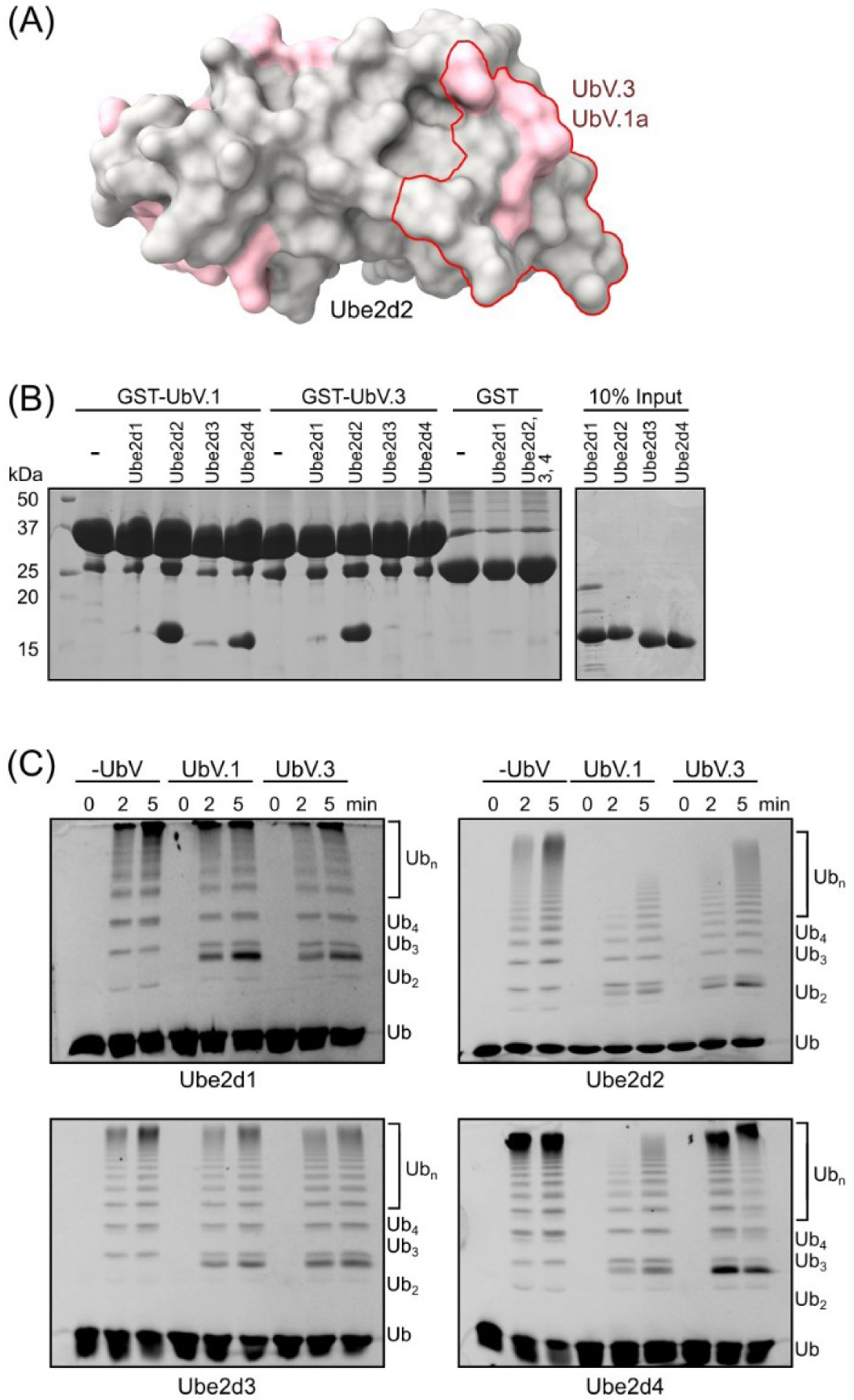
Specificity of the UhVs for the Ube2d family is high. **(A)** A surface representation of Ube2d2 with amino acids that vary between the four Ube2d family members highlighted in pink. The interface with UbV.la and UbV.3 is outlined in red. UbV.lb does not contact varied Ube2d residues. **(B)** GST pulldown experiments comparing the binding of GST-UbVs to the four members of the Ube2d family of E2 enzymes. **(C)** Multi-turnover ubiquitin chain building assays with the four Ube2d family members in the presence or absence of the UbVs. Ubiquitin monomers and polyubiquitin were visualised by fluorescence of Cy3 at 600 nm.

## Discussion

Attachment of the correct type of ubiquitin to substrate proteins is highly dependent on the enzymes involved. While the E3 ligases are the most diverse and bind specific substrate proteins, the E2 enzymes play an essential role in deciding the exact kind of ubiquitin chain to be attached. Because the downstream consequence for the substrate is dependent on the exact nature of the ubiquitin code, it is important to understand how E2 enzymes work and are regulated. Here, we used phage display to discover UbVs that are capable of binding to and specifically disrupting E2 enzyme function. Two of the UbVs reduced engagement with the E1 enzyme, resulting in a decrease in the amount of charged E2 enzyme and a consequent decrease in its activity. Along with blocking the E2:E1 interaction, the more inhibitory UbV, UbV.1 also likely sterically inhibited allosteric backside ubiquitin binding (Fig 3D), and this may account for its greater ability to impede chain-building.

Other compounds have been discovered that bind to many E2 enzymes [1], and there are some reports of proteins and small molecules that can modulate the activity of the Ube2d family. For protein-based inhibitors, a UbV isolated using similar technology as reported here could specifically inhibit Ube2d1 by directly blocking backside ubiquitin binding [17]. Whereas at least three sesquiterpene lactones and a triterpenoid derived from medicinal herbs were reported to inhibit the activity of Ube2d [22–25]. These small molecules are either broadly specific for the Ube2d family or specificity was not reported. More recently, two small molecules were discovered that bind at an allosteric site on the Ube2d family [26,27]. One of these can act as a molecular glue to promote degradation of NFκβ1, and both small molecules were developed into Proteolysis Targeting Chimeras (PROTACs) and shown to degrade model substrates. However, while chemoproteomic profiles suggest that the small molecules are specific to the Ube2d family, no differentiation could be made between members of the Ube2d family.

The two UbVs characterised in detail here both appear to inhibit interaction between Ube2d2 and the E1 enzyme (Fig 3A), and because of the sequence similarity between all the UbVs reported here (Fig 1A), it is likely that the other four bind at the same interface. When bound to Ube2d2, the UbVs are predicted to sterically inhibit interactions with the ubiquitin fold domain (UFD) of the E1 enzyme, and this is supported by the slowed charging of Ube2d2 in the presence of UbV.1 and UbV.3. The affinity between the E1 enzyme and Ube2d has been reported to range from a K_m_ of 0.2 to 10 µM [28,29], and the UbVs reported here bind at the higher end of this scale. This suggests that the UbVs are able to compete with the E1 enzyme for binding to the E2. It is possible that improving the binding affinity of the UbVs for Ube2d2 by affinity maturation would result in discovery of UbVs that are more effective inhibitors. Interestingly, this same E1-E2 interface was also blocked by two recently discovered UbVs that bind another E2 enzyme, Ube2k [18]. This finding suggests that the surface of E2 enzymes comprising the α1 helix and the β1-β2 loop may be a hotspot that could be targeted in an E2-specific way to disrupt ubiquitin transfer activity.

One of the strengths of phage display is its ability to select for interactions that are highly specific. Discovery of highly-specific binders using phage display has been seen before, including between members of highly-similar proteins [14,30,31]. Similarly, the UbVs we have discovered are specific for particular members of the Ube2d family, and at least one appears to bind only two members of the Ube2d family to the exclusion of all other tested E2 enzymes. There are few tools that can distinguish activity within closely related proteins, and this has resulted in a possibly inaccurate assumption that highly similar family members play redundant roles in cells. This feature makes the UbVs reported here, and elsewhere, potentially immediately useful as research tools to help investigate the biological role of specific members of the Ube2d E2 enzyme family.

## Materials

### Cloning and protein expression and purification

For phage display, Ube2d2^S22R^ was cloned into pET-LIC including a cleavable His-tag and a 3’ Avi tag for biotinylation. The E2 enzymes and RNF12^RING^ were cloned into a pGex6P3 vector, as previously described [32]. Ubiquitin was cloned into a pET3a vector, and the E1 enzyme was cloned into a pET24a vector as previously described [33,34]. The UbVs were amplified from the phagemid vector with primers that included 5’ and 3’ ligation independent cloning sticky ends and cloned into both a pET-LIC and a pET-pGex vector. The amplified UbV constructs included a 5’ sequence that encoded a FLAG tag.

All recombinant protein production was performed in *E. coli* BL21 (DE3) cells grown in LB media. Each culture was grown at 37 °C until the OD_600_ was 0.6, then protein expression was induced by the addition of 0.2 mM IPTG and the cells were incubated at 18 °C for 16 h. The bacterial pellet was harvested, resuspended in PBS, and sonicated (Sonifier Cell Disrupter, Branson) before being centrifuged (Avanti J-26 XP) in an F50C rotor for 25 min at 12,000 rpm to remove cell debris. The E2 enzymes, Ube2d2 and Ube2d2^S22R^ were expressed with a cleavable N-terminal GST tag from the pGEX-6P-3 vector. After centrifugation of cell lysis, the supernatant was mixed with GSH resin (GE Healthcare) and left to rotate for 1 hr at 5 °C. The GSH resin was washed, then the protein was digested using in-house produced 3C to cleave the GST tag. A final size-exclusion run (Superdex 75 column, Cytiva) was performed. The UbVs and Ube2d2^S22R^-Avi were purified by nickel-affinity chromatography with HIS-select resin (Sigma) followed by an overnight 3C digestion using in-house produced 3C protease. Subsequently proteins were purified with a Superdex 75 column (Cytiva) equilibrated in PBS. Elutions of all proteins were pooled and concentrated before being flash frozen in liquid nitrogen and stored at −80 °C. Ube2d2S22R-Avi was biotinylated using in-house produced BirA as previously described [18]. The E1, RNF12^RING^, and ubiquitin were purified as previously described [32–35].

### Phage display of UbVs binding to Ube2d2^S22R^

Phage display was performed as described [17,18]. In short, biotinylated Ube2d2^S22R^ was immobilised onto Nunc MaxiSorp 96-well plates (Thermo Fisher Scientific) that were coated with streptavidin or neutravidin (New England BioLabs). Four rounds of phage display binding selection were performed. Ninety-six clones were screened for binding to Ube2d2 using ELISA before positive clones were sequenced. From these sequences, six UbVs were selected and investigated.

### In vitro ubiquitin assays

For chain-building assays, 0.1 μM E1, 5 μM Ube2d2, 50 μM ubiquitin, 10 μM Cy3-labelled ubiquitin, 7 μM RNF12^RING^ were mixed with or without 50 μM UbVs in an assay buffer (50 mM Tris–HCl pH 7.5, 50 mM NaCl, 2 mM MgCl_2_, 2 mM TCEP and 5 mM ATP) and incubated at 37°C. The reaction was quenched at the indicated timepoints by mixing with reducing SDS-PAGE buffer and samples were resolved using SDS-PAGE. Labelled ubiquitin was visualised using an Odyssey Fc system (LI-COR) at 600 nm with a 2 min exposure. For ubiquitin charging assays, 0.1 µM E1, 20 µM Ube2d2, 30 µM ubiquitin, and 10 µM Cy3-labelled ubiquitin were mixed with or without 200 μM UbVs in assay buffer and incubated at 28°C for the indicated time points. Samples were resolved using SDS-PAGE and visualised at 600 nm. Ubiquitin discharge assays were performed with Ube2d2∼Ub that was produced with Cy3-labelled ubiquitin then purified with an S75 Superdex column. For the discharge experiment, 5 µM of Ube2d2∼Cy3-ubiquitin, 2 µM of RNF12^RING^, and 5 µM lysine were mixed with or without 50 µM UbVs and incubated at 25 °C. At the indicated timepoints, samples were removed and the reaction was quenched with non-reducing SDS-PAGE loading buffer. SDS-PAGE gels were imaged at 600 nm. Quantification of remaining conjugate was performed in technical triplicate with Image Studio Lite (LI-COR Biosciences).

### Enzyme-linked immunosorbent assays

For the specificity ELISA experiments, a representative library of E2 enzymes was prepared, as described elsewhere [18]. These were immobilised to 96-well Maxisorp plates (Thermo Fisher) and probed with FLAG-tagged purified UbVs. After binding of the UbVs for 1 h, HRP-tagged FLAG antibody was added to the wells, before mixing with TMB substrate for colourimetric visualisation of antibody interactions.

### Crystallography and structure solution

UbV-Ube2d2 complexes were produced by mixing UbVs at a 2-fold molar excess with Ube2d2 or Ube2d2^S22R^. The samples were incubated on ice for 60 min then the complexes were purified using a 10/300 S75 (Cytiva). Subsequently the complexes were concentrated to 5-10 mg/mL and mixed at a 1:1 and a 2:1 v/v ratio with PACT, JCSG, and ProPlex crystal screens (Molecular Dimensions) with a mosquito (TTP Labtech). The UbV.1-Ube2d2 complex grew diffraction-quality crystals in 0.1 M Sodium citrate pH 5.6, 20% w/v PEG 4000, and 20% v/v 2-propanol. The UbV.3-Ube2d2^S22R^ complex crystallised in 0.1 M Sodium HEPES pH 7.0, and 15% w/v PEG 20,000. Data from these crystals were collected at the Australian Synchrotron on beamline MX2 and processed with XDS [36], merged with Aimless, and their structures solved using Phaser [37,38]. Subsequent refinement was performed in Phenix Refine [39] followed by iterative modelling in coot v0.9.8.3 [40]. All structural images were produced with ChimeraX v1.6.1 [41].

### Binding experiments

Isothermal titration calorimetry (ITC) experiments were performed in a MicroCal VP-ITC at 25 °C. For each ITC experiment, Ube2d2 or Ube2d2^S22R^ at approximately 20 µM was in the sample cell while the UbVs or ubiquitin (approximately 200 µM) were in the syringe. All samples were either purified in or desalted into a common stock of PBS. Data was processed with NITPIC, SEDPHAT, and GUSSI [42]. For the pulldown experiments, GST-UbVs, GST-RNF12^RING^ and the GST control were purified by binding to GSH resin and washing resin extensively. The resin-bound samples were mixed with the candidate proteins and incubated at 4 °C for 1 hour before being washed in triplicate and resolved with SDS-PAGE.

### Analytical size exclusion chromatography

Analytical size exclusion chromatography was performed using a Superdex S75 10/300 column (Cytiva) equilibrated with PBS. Single proteins were injected at a final concentration of 50 µM while for mixtures, Ube2d2 and Ube2d2^S22R^ at 50 µM were mixed with UbVs at 100 µM and incubated on ice for approximately 30 min. The column was calibrated with ribonuclease A (13.7 kDa), carbonic anhydrase (29 kDa) and ovalbumin (43 kDa) to estimate the molecular weight of eluted complexes.

### Analytical ultracentrifugation

Sedimentation velocity experiments were performed using a Beckman Coulter Optima analytical ultracentrifuge. For the experiments to determine the characteristics of individual proteins, UbV.1 (0.5 mg/mL, 50 µM), UbV.3 (0.5 mg/mL, 50 µM), Ube2d2 (0.25 mg/mL, 15 µM), and Ube2d2^S22R^ (0.25 mg/mL, 15 µM) were each detected at 280 nm. Peaks in the observed distributions that could represent larger species (at ∼2.1 S for UbV.1, UbV.3 and at ∼3.1 S for Ube2d2, Ube2d2^S22R^) represented less than 5% of the signal and were not further investigated. Ube2d2 and Ube2d2^S22R^ were fluorescently labelled with FITC following the manufacturer instructions provided in the FITC labelling kit (Thermo Fisher Scientific). For the analysis of UbV-Ube2d2-FITC complexes, the UbVs (1.0 mg/mL 100 µM) were mixed with Ube2d2-FITC and Ube2d2^S22R^-FITC (0.5 mg/mL 30 µM) and FITC fluorescence was measured at 493 nm. All data were collected at 25 °C in PBS pH 7.4 in 12 mm double sector cells with sapphire or quartz windows and were run in an An-50 Ti rotor at 50,000 rpm. Sedimentation data were analysed with UltraScan 4.0 [43,44]. Optimisation was performed by two-dimensional spectrum analysis (2DSA) [45,46] with simultaneous removal of time- and radially invariant noise contributions and fitting of boundary conditions. The 2DSA solutions were subjected to parsimonious regularisation by genetic algorithm analysis [47].

## Acknowledgements

This research was supported by a Sir Charles Hercus Health Research Fellowship administered by the Health Research Council of new Zealand awarded to AJM. This research was undertaken in part using the MX2 beamline at the Australian Synchrotron, part of ANSTO, and made use of the Australian Cancer Research Foundation (ACRF) detector. We also thank the New Zealand Synchrotron Group for facilitating access to the Australian Synchrotron.

## Author contributions

**Jeffrey McAlpine:** investigation (equal), formal analysis, visualisation (equal). **Jingyi Zhu:** investigation (equal), visualisation (equal). **Nicholas Pudjihartono:** investigation. **Joan Teyra:** methodology, writing – review & editing. **Michael Currie:** investigation, formal analysis. **Renwick Dobson:** formal analysis, software, resources, writing – review & editing. **Sachdev Sidhu:** methodology, resources, writing – review & editing. **Catherine Day:** writing – review & editing, supervision (supporting), resources. **Adam Middleton:** conceptualisation, formal analysis, funding acquisition, investigation, project administration, supervision, writing – original draft preparation, writing – review & editing.

## Supplementary figures

**Supplemental Figure 1.**
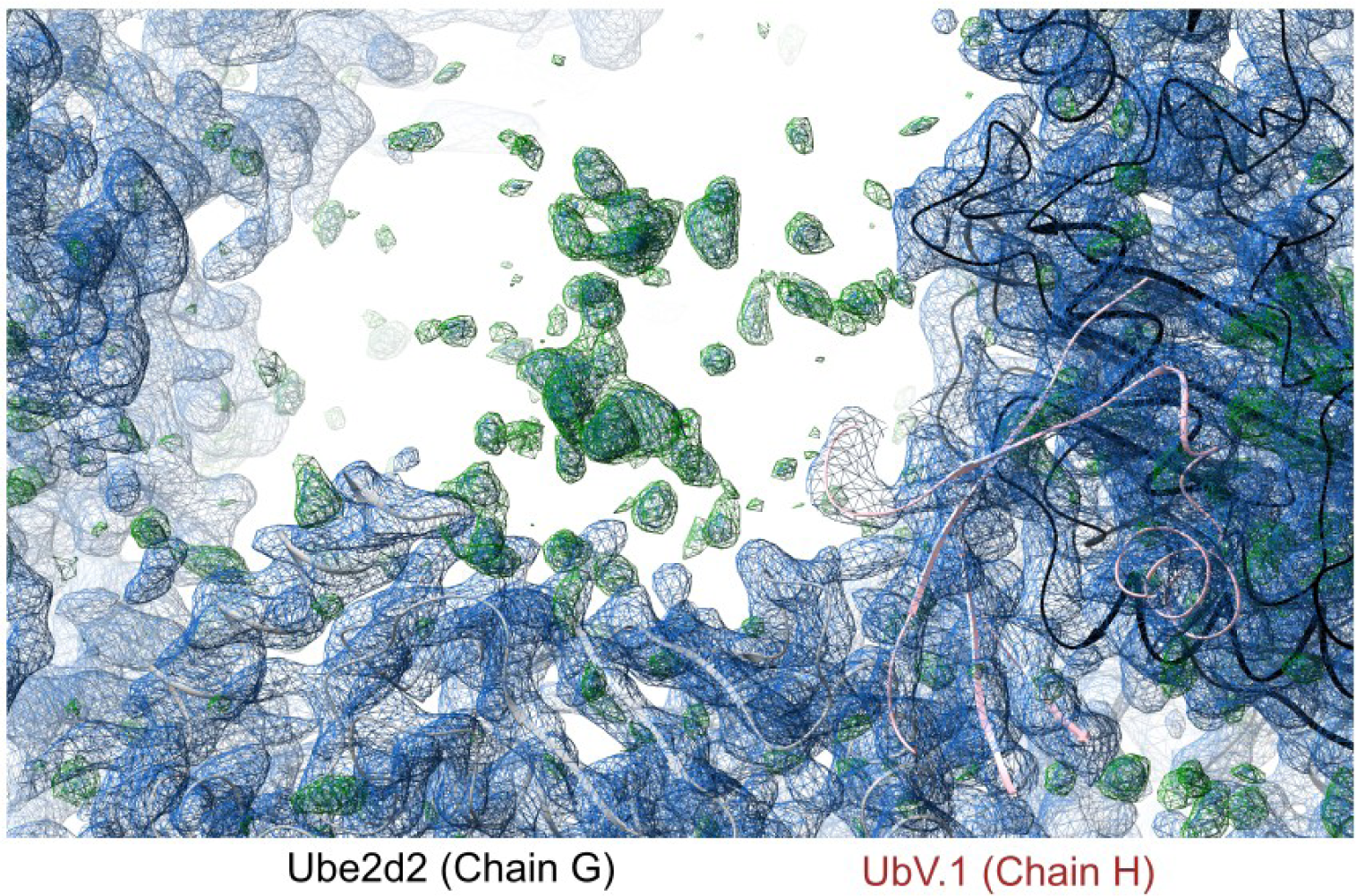
Structural characterisation of the UbV.1-Ube2d2 complex. Electron density map showing where UbV.1b could not be modelled against the Chain G model of Ube2d2. In blue mesh is the 2mFo-DFc shown at an RMSD of 1.2 Å while green mesh shows the mFo-DFc at an RMSD of 2.5 Å.

**Supplementary Figure 2.**
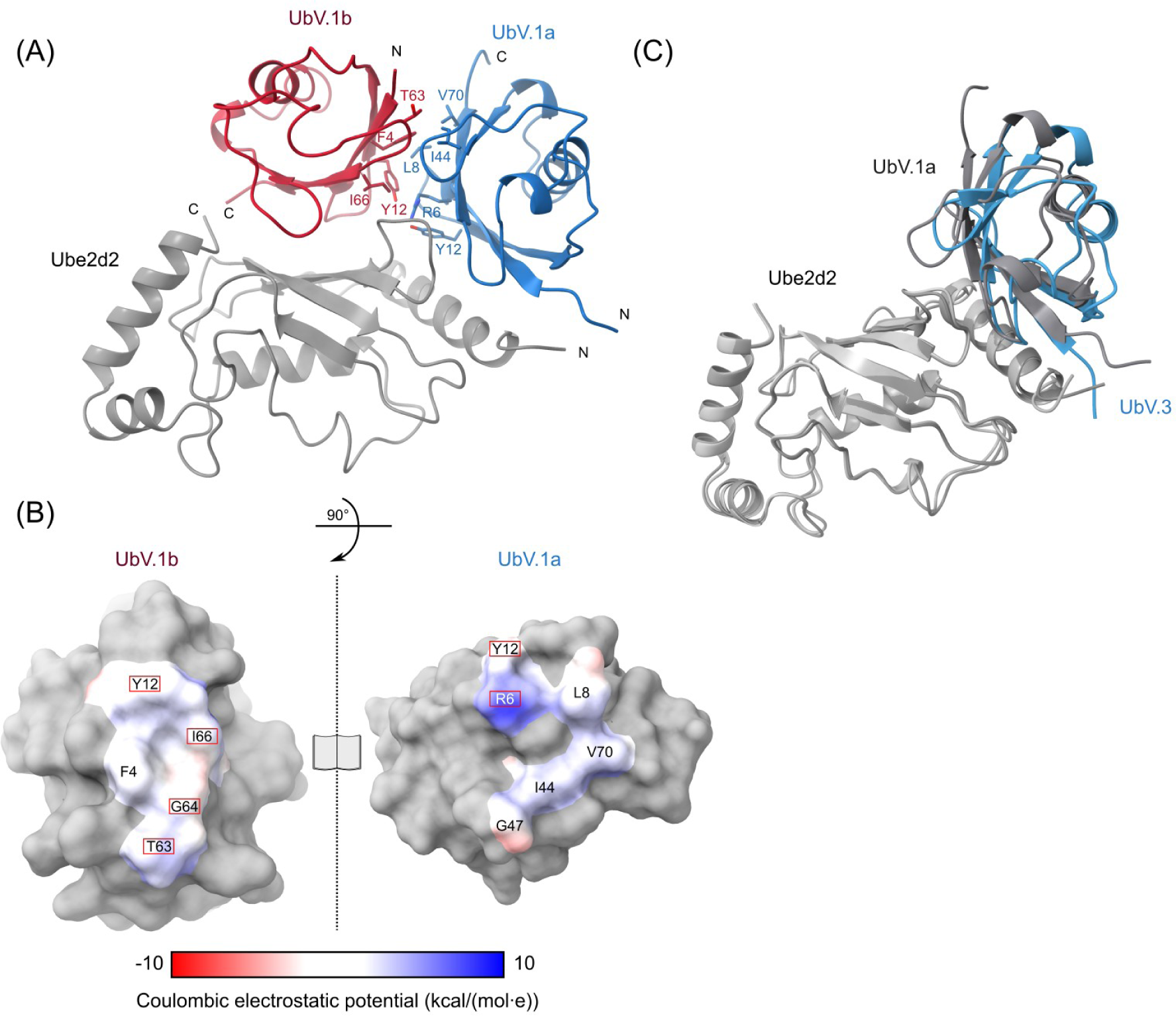
Analysis and comparison of the binding of the two UbVs. **(A)** Contacting residues between the two molecules of UbV.1 shown as sticks. **(B)** Open-book surface representation showing the contacting residues between two UbV.1 molecules. Colouring is based off the Coulombic electrostatic potential of the proteins as calculated by Chimerax v1.6.1 [41]. Residues that are diversified from wild-type ubiquitin are highlighted with red boxes. **(C)** An overlay of Ube2d2-UbV.1a with Ube2d2^S22R^-UbV.3 to compare the orientation of the UbVs. This highlights the approximate 10° rotation between the two UbV molecules.

**Supplementary Figure 3.**
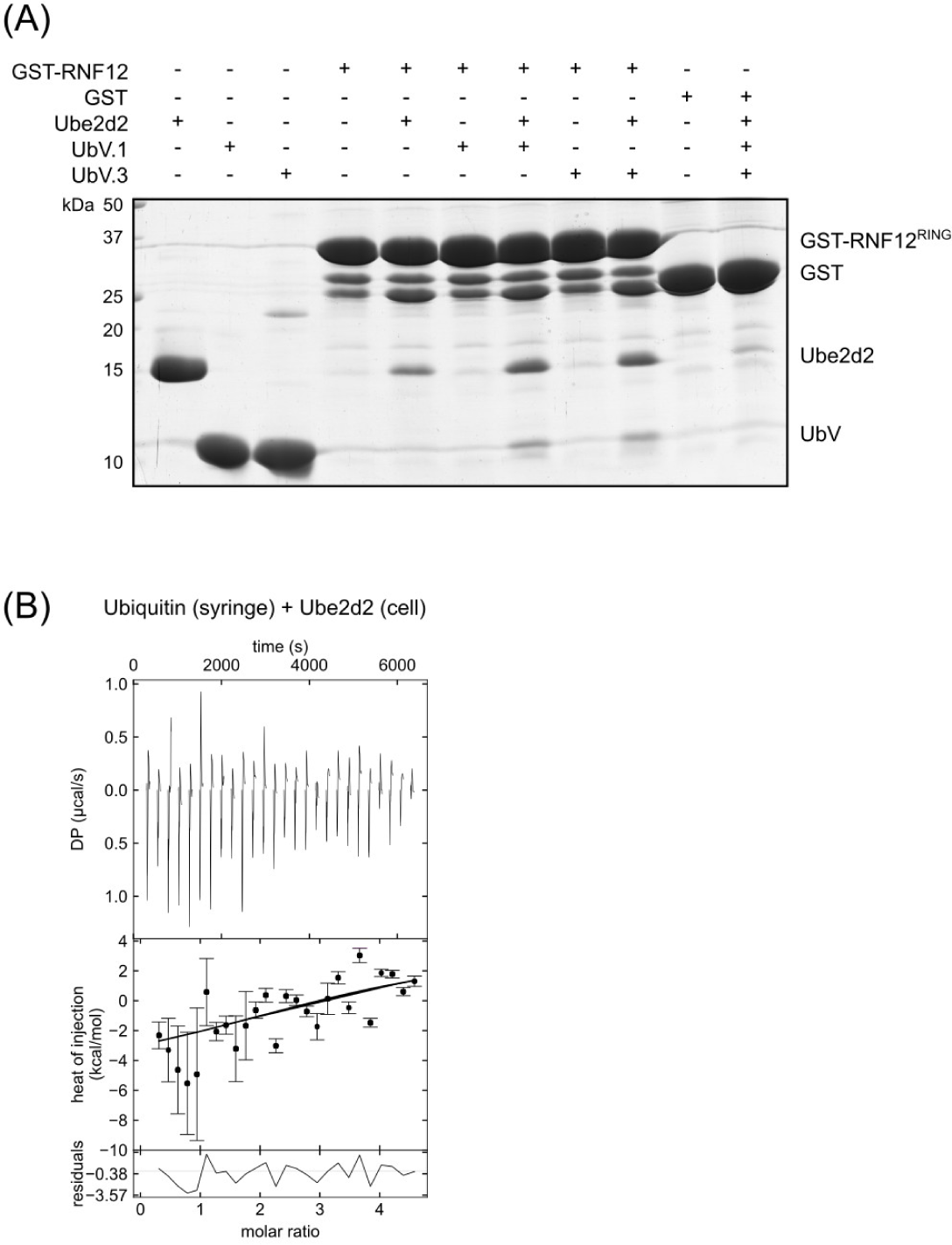
Analysis of interaction between UbVs and ubiquitin with Ube2d2. **(A)** A pulldown experiment comparing binding of GST-RNF12^RING^ to Ube2d2 with or without the UbVs. **(B)** ITC where wild-type ubiquitin (200 µM) was titrated into Ube2d2 (20 µM).

**Supplementary Figure 4.**
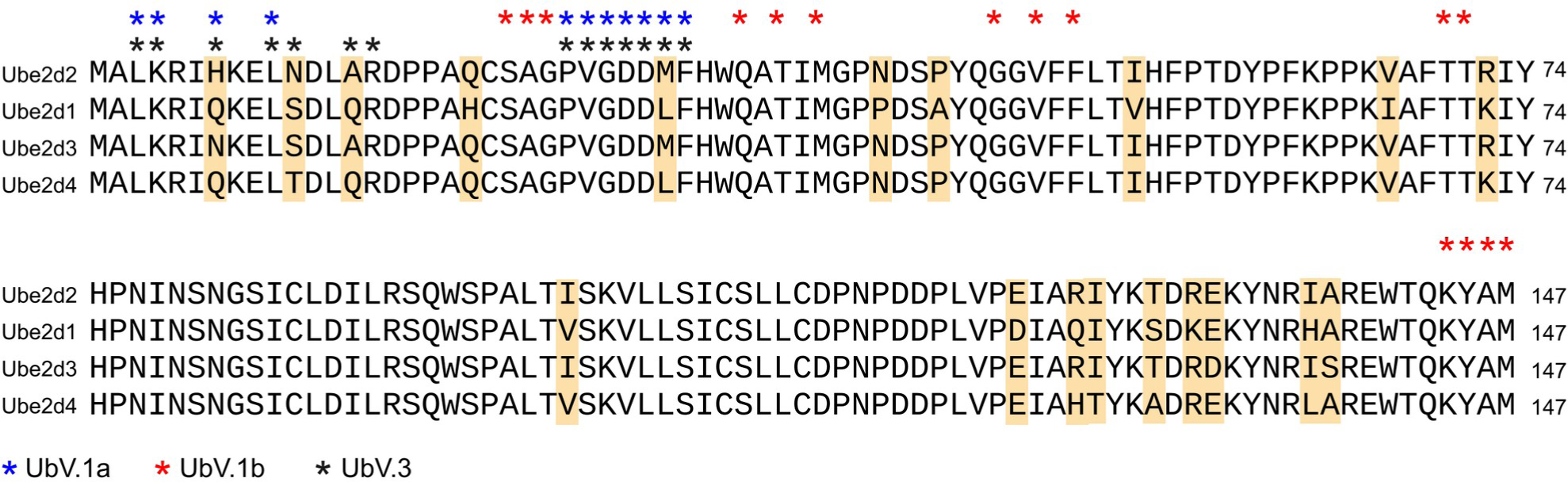
Conservation within the Ube2d family of E2 enzymes. Amino acid alignment of the four members of the Ube2d family. Highlighted in beige are variable residues in the family. Coloured asterisks indicate residues that interact with the UbVs.

